# The potato cyst nematode *Globodera pallida* possesses an expanded serotonergic nervous system

**DOI:** 10.64898/2026.09.29.755464

**Authors:** Mirela C. Coke, Catherine J. Lilley, Roberta Healey, Vincent O’Connor, Lindy Holden-Dye, Peter E. Urwin

## Abstract

The neurotransmitter serotonin (5-hydroxytryptamine; 5-HT) controls behaviours that are essential for the parasitic lifestyle of plant parasitic nematodes (PPNs). Here, we used serotonin immunochemistry and *in situ* hybridization for the serotonin synthesis genes, *Gpa*-*tph-1* and *Gpa*-*bas-1*, to map its signalling pathways in *Globodera pallida* J2 infective stages. We identified fourteen serotonin-immunoreactive cell bodies around the pharyngeal median bulb and revealed that eleven of them co-express *Gpa*-*tph-1* and *Gpa-bas-1,* confirming these are serotonin-synthesising neurons. Serotonin immunoreactivity was also localised in structures consistent with labelling of amphids, and in sixteen non-neuronal cells located in the middle and posterior region of the nematode. Depletion of serotonin with reserpine reduced the serotonin immunoreactivity in all fourteen cell bodies confirming their neuronal identity, but not in non-neuronal tissue. We identified additional, but infrequent expression of *Gpa*-*tph-1* in two cells and expression of *Gpa*-*bas-1* in one pair of symmetrically bilateral cells anterior to the median bulb, that lacked serotonin immunoreactivity. The results reveal distinct pools of serotonin distribution in *G. pallida* and indicate serotonin signalling is more complex than in other parasitic nematodes and *C. elegans*, highlighting evolutionary divergence in neurotransmitter identity.

## Introduction

The cyst nematode *Globodera pallida* is a sedentary endoparasite of potato that causes significant economic losses worldwide (Cotton et al., 2014). In the absence of a single dominant resistance gene to support the development of resistant crops and with a slow annual decline rate of dormant *G. pallida* populations in the soil that makes crop rotation economically inefficient, the control of these nematodes is achieved mainly by chemical means (Trudgill et al., 2014, Crisford et al., 2020). The majority of current nematicides target the neuromuscular system of the nematode or are metabolic inhibitors (Holden-Dye and Walker, 2014, Kearn et al., 2014, Schleker et al., 2022). However, several nematicides have been withdrawn from use due to elevated levels of ecotoxicity while reports suggest that others are losing their efficacy (Desaeger et al., 2020). Replacing the withdrawn nematicides is a slow process and sparse knowledge of the nervous system of plant parasitic nematodes compounds the difficulties with developing new control strategies. The most pressing knowledge gap in this regard is a lack of a detailed understanding of the neural and physiological basis of the behaviours that are essential for a successful parasitic life-cycle (Coke et al., 2024). *Globodera pallida* hatches as motile second-stage juveniles (J2s) from encysted eggs in the soil only in response to chemicals released from the roots of a host plant (Lilley et al., 2005). The hatched J2s move towards the host roots and use their anterior stylet, a spear-like organ, to pierce the plant cell wall. Once inside the host root, they migrate intracellularly in a destructive way towards the vascular cylinder where their behaviour changes as they gently probe for a suitable cell that once selected will become the initial feeding cell (Sobczak and Golinowski, 2011). The nematode delivers pharyngeal secretions through the stylet into this initial feeding cell, reprograming the plant cells into a specialized feeding site that supports development of the nematode until the end of its life cycle. These behaviours are tightly under neuronal control and represent important targets for the development of new control strategies (Han and Schroeder, 2025).

Serotonin (5-hydroxytryptamine, 5-HT) is a key neurotransmitter controlling behaviours that are essential for the early interaction of the J2 with the plant root. Activation of stylet thrusting is critically dependent on serotonin (Crisford et al., 2020, Masler, 2008). Therefore, disruption of serotonin signalling in J2 is an attractive focus for research. In line with this, reserpine, a plant alkaloid from the Indian snakeroot has been found to act as a crop protectant against *G. pallida*. Reserpine inhibits the vesicular monoamine transporter, VMAT, which is responsible for loading serotonin into synaptic vesicles for neuronal release (Erickson et al., 1992), thereby inhibiting serotonin signalling. In *G. pallida* reserpine depletes neuronal serotonin and this prevents activation of stylet thrusting thus impairing the nematode’s ability to access the plant root (Crisford et al., 2020). Additionally, the serotonin receptor MOD-1, which appears to be restricted to phylum Nematoda, has been suggested as a potential target for developing future control strategies (Rodriguez Araujo et al., 2022, Calahorro et al., 2022). As the interest around serotonin increases, core components of the serotonergic signalling pathway in *G. pallida* have been functionally characterized and their identity confirmed by successful complementation of their corresponding *C. elegans* mutants (Crisford et al., 2020). However, despite the molecular data on the presence and function of endogenous serotonin signalling components, there is no information regarding serotonin distribution or the identity of serotonergic neurons in the nervous system of *G. pallida*. This contrasts with the extensive neurobiological studies and the detailed neuronal and neurotransmitter maps available for the free-living nematode *Caenorhabditis elegans* (Yemini et al., 2021, Wang et al., 2024). The neuronal localisation of serotonin is well known in *C. elegan*s along with the distinction between serotonin-synthesising neurons and the serotonin-uptake neurons (Horvitz et al., 1982, Sawin et al., 2000).

Neurons that synthesise serotonin are defined by the presence of the rate-limiting enzyme, tryptophan hydroxylase (TPH) encoded by the gene *tph-1* (Sze et al., 2000). In addition, they must express the aromatic amino acid decarboxylase BAS (Hare and Loer, 2004) thus enabling hydroxylation of the substrate, tryptophan, and its subsequent decarboxylation to serotonin. These criteria define the presence of a small set of serotonergic neurons, NSM, ADF and HSN in the *C. elegans* hermaphrodite (Sze et al., 2000). Additionally, serotonin is synthesized in the CP and tail neurons in the male *C. elegans* (Loer and Kenyon, 1993). Some neurons and tissues may accumulate serotonin from their surroundings via the plasma membrane transporter MOD-5 in the absence of TPH-1 (Jafari et al., 2011). This is the case for *C. elegans* AIM and RIH neurons (Serrano-Saiz et al., 2017, Jafari et al., 2011). Weak accumulation of serotonin has also been reported in the URX and I5 neurons in the head of both sexes, in the VC04/VC05 ventral cord motor neurons in the hermaphrodite and the CEM neurons of males only (Sawin et al., 2000, Serrano-Saiz et al., 2017, Jafari et al., 2011).

Neuronal architecture is considered to be well conserved in the phylum Nematoda (Hong et al., 2019) and the few existing studies in plant parasitic nematodes support this idea (Kimber et al., 2002). Differences in the serotonin distribution in the nervous system of free-living nematodes have been reported, however no link between behavioural outputs and these particular differences has been identified as yet (Rivard et al., 2010). The nervous systems of the few plant parasitic nematodes analysed showed a similar pattern to that of *C. elegans,* with serotonin immunolocalized in putative homologues of ADF, NSM, VC and CP neurons in the lesion nematode *Pratylenchus penetrans* (Han et al., 2017). In J2s of the root-knot nematode *Meloidogyne graminicola* serotonin immunoreactivity labelled cell bodies that may be homologues of ADF, NSM and RIH (Yang et al., 2024). In both species, one pair of putative cell bodies located anterior to the nerve ring and lacking a *C. elegans* counterpart, was serotonin-immunoreactive (IR).

Here, the serotonin distribution in the nervous system of *G. pallida* J2s was investigated, along with the mapping of serotonergic neurons, to provide the most complete description of a serotonergic system to date for a plant parasitic nematode. We identified a unique serotonin distribution pattern for *G. pallida*, distinct from other plant parasitic nematodes, characterized by an expanded number of serotonin-synthesising neurons. The results suggest evolutionary plasticity in the serotonergic system marked by a potential shift in neuronal identity for putative homologous neurons.

## Material and Methods

### Nematode culture

The soil-extracted cysts of *G. pallida* population Lindley (Pa2/3) were exposed to fresh potato root diffusate to stimulate hatching as described previously (Crisford et al., 2020). Hatched J2s were collected daily over a week, washed twice with tap water then pooled and stored at 10 °C prior to use.

### Reserpine treatment

Reserpine (Sigma-Aldrich) was dissolved at 30 mM in dimethylformamide (DMF) and stored in aliquots at –20 °C. J2s were soaked in 100 µM reserpine diluted in filter-sterilized tap water or in 0.33% DMF for the vehicle control, for 4, 8 and 24 hours prior to fixation and immuno-localisation.

### Serotonin immuno-localisation

*G. pallida* J2s were fixed, sectioned and permeabilized as described previously (Thorpe et al., 2014). The processed J2s were blocked for 30 minutes (min) in 1% blocking reagent (Roche) prior to incubation overnight, at 4°C, in 1:200 rabbit-derived anti-serotonin primary antibody (Sigma-Aldrich). The nematodes were washed and re-blocked for 30 min in 1% blocking reagent (Roche) prior to incubation for 2 hours at room temperature in 1:200 goat-derived FITC-labelled anti-rabbit secondary antibody. For control experiments the J2s were treated in the same manner as above except the addition of the primary antibody was omitted from the protocol. Samples were washed and resuspended in a few drops of Citifluor anti-fadent. Images were acquired with either a Leica DMRB microscope fitted with a QIcam camera and connected to Q-Capture software, or with a LSM 800 confocal microscope connected to ZEN software.

### Phalloidin staining

Phalloidin staining of muscles was performed following serotonin immunostaining. The immunostained J2 nematodes were incubated overnight at 4°C in 200 ng/ml Phalloidin-tetramethylrhodamine conjugate (Santa Cruz Biotechnology) and washed with PBS before resuspension in a few drops of anti-fadent. Images were acquired as described above.

### In situ hybridisation using HCR probes

All sets of probes used were designed by Molecular Instruments based on the coding sequences of GPLIN_000790300 (*Gpa-tph-1*) and GPLIN_000551500 (*Gpa-bas-1*). For *Gpa-tph-1*, one set of 14 probes was designed from the first 754 bases and was used with the B1 amplifier labelled with Alexa 488 fluorophore. The same set was also used with the B2 amplifier labelled with Alexa 647 fluorophore. A second set of 14 probes designed from the last 845 bases was used with the B2 amplifier labelled with Alexa 647 fluorophore. For *Gpa-bas-1*, one set of 20 probes was used with the B3 amplifier labelled with Alexa 546 fluorophore. The full set of 28 probes for *Gpa-tph-1*, all using the B2 amplifier labelled with Alexa 647, was used for multiplexed co-detection of *Gpa-tph-1* and *Gpa-bas-1*.

*G. pallida* J2s were fixed, sectioned and permeabilized as described above. The subsequent *in situ* hybridization followed the protocol described previously (Sperling and Eves-van den Akker, 2023, Choi et al., 2016). The J2s were washed for 5 min in 1:1 Hybridization Buffer (Molecular Instruments) and 1% PBST (PBS with 1% Tween). The nematodes were pre-hybridized in Hybridization Buffer (Molecular Instruments) for 30 min prior to incubation overnight in hybridization buffer with 1% probe stock solution. The J2s were washed three times, 15 min each time, in 37°C pre-warmed Probe Wash Buffer (Molecular Instruments) and once for 5 min in 5xSSCT with 0.1% Tween prior to incubation for 30 min in amplification buffer. The nematodes were then incubated for 20-22 hours at room temperature with the amplification hairpins prepared as detailed in Choi et al. (2016). The nematodes were washed with 5xSSC containing 0.1% Tween prior to re-suspension in mounting medium (Slow Fade Gold antifade mountant). Images were acquired as described above.

### DAPI staining

Following immuno-localisation and fluorescence ISH J2s were incubated for 30 min in 0.5 µg/ml DAPI (Sigma-Aldrich). The J2s were then washed three times with water prior to observation.

## Results

### Serotonin localises to multiple candidate neurons and the amphids in J2s of *G. pallida*

Immunolocalization was used to determine the distribution of endogenous serotonin in *G. pallida* J2s. Putative serotonin-IR cells were identified in the anterior, middle and posterior region of the body (Fig. 1, Fig. 2). We consider the head and the pharyngeal area to be the anterior region of the body. The two large cells of the genital primordium could be reliably identified at approximatively 60% distance from the head of the nematode and were used as a landmark (Fig. 2). We consider the region anterior to the genital primordium but posterior to the cell bodies near the median bulb as the middle region of the nematode. The region between the genital primordium and the tail of the nematode will be referred to as the posterior region. Two pores at the head of the J2 along with two symmetrically bilateral structures consistent with the position of the amphid pouches displayed strong fluorescence (Fig. 1A). Phalloidin staining of muscles confirmed that the fluorescence did not correspond to the position of the stylet protractor muscles (Supp. Fig. 1). Two sets of symmetrically bilateral putative cell bodies we have named A1,2 and B1,2 for clarity can be distinguished in the pharyngeal median bulb (Fig. 1B). The A1,2 pair, with moderate serotonin immunoreactivity is located anterior to the median bulb valve. These putative cell bodies do not have a positional counterpart in *C. elegans* or other plant-parasitic nematodes studied to date. However a similar location for a pair of serotonin-IR putative cell bodies was reported in the free-living nematodes *Oscheius myriophila, Pristionchus pacificus* and *Panagrellus redivivus* (Rivard et al., 2010). The B1,2 pair of putative cell bodies, located posterior to the median bulb valve, is consistently (90%, n=10) and strongly stained. Their position suggests they might be homologues of the *C. elegans* NSM neurons (Horvitz et al., 1982). Another two pairs of symmetrically bilateral putative cell bodies, D1,2 and E1,2, can be distinguished posterior to the median bulb (Fig. 1B). The pair D1,2 displayed faint serotonin immunoreactivity. Putative cell bodies in similar position have been reported in both *P. penetrans* and *M. graminicola* (Han et al., 2017, Yang et al., 2024). Considering their position, they could be homologues of the *C. elegans* URX neurons which only uptake serotonin through an unknown mechanism (Serrano-Saiz et al., 2017). The pair E1,2 are putative homologues of the ADF neurons from *C. elegans* (Sze et al., 2000) and consistently (90%, n=10) displayed a strong fluorescence. At least one unpaired cell body, C (Fig. 1B), can be identified in between these pairs of cell bodies, with no counterpart in *C. elegans* or other plant-parasitic nematodes.

**Figure 1.**
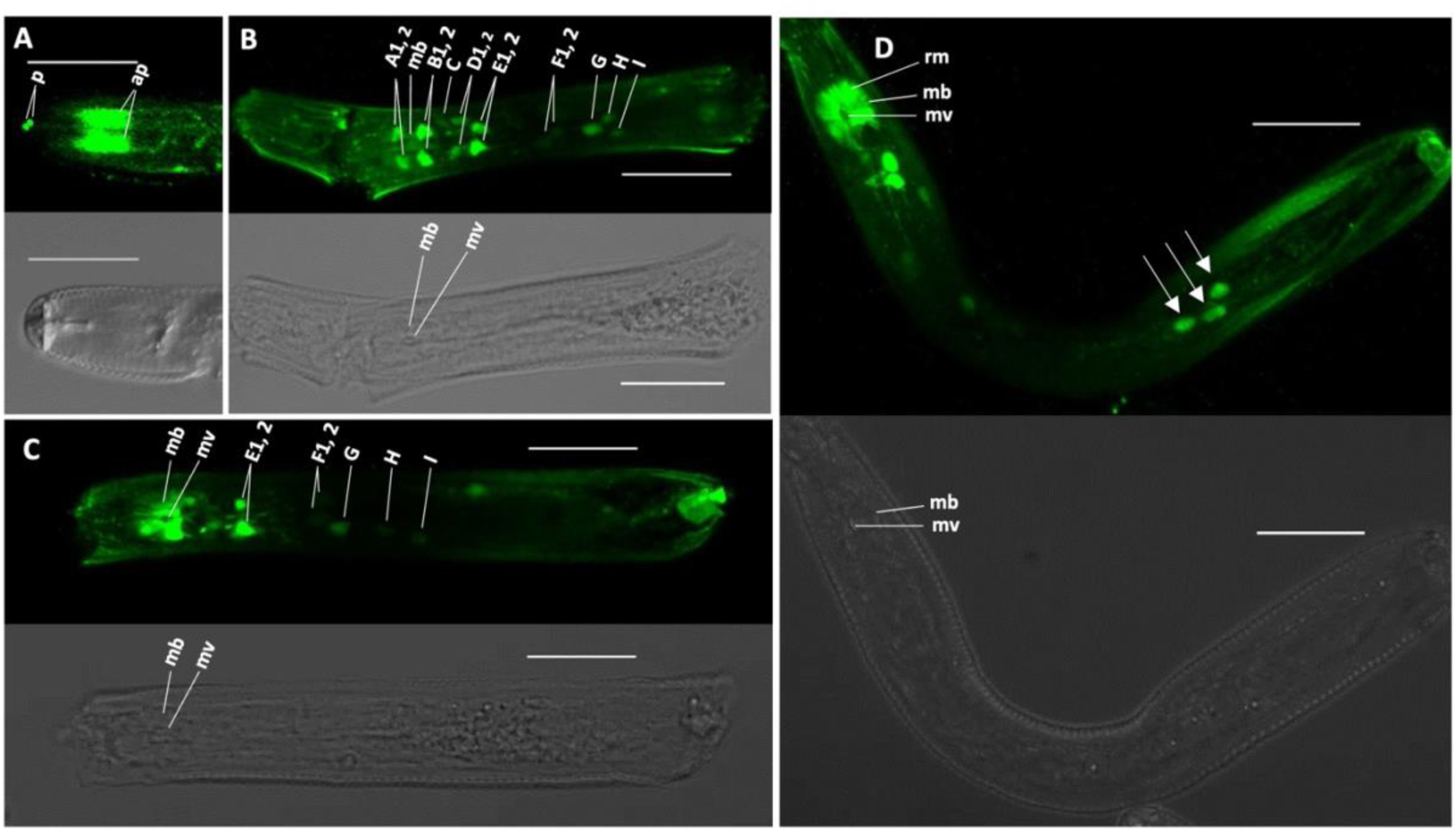
Serotonin immunolocalization in 2nd-stage juveniles (J2s) of *G. pallida*. Anterior region of the *G. pallida* J2 is to the left in all images. **(A)** Strong anti-serotonin staining is present in two pores (p) and two structures resembling the amphid pouches (ap) in the head of the nematode. **(B)** Staining can be detected in fourteen cell bodies (A1,2, B1,2, C, D1,2, E1,2, F1,2, G, H, I) in or adjacent to the pharyngeal median bulb (mb). Strong staining was consistently detected in two symmetrically bilateral pairs of cell body - B1,2 and E1,2 and their position suggest they could be putative homologues of NSM and ADF neurons respectively. **(C & D)** Occasionally, strong staining can be detected in the median bulb (mb) radial muscles (rm), excluding the median bulb valve. **(D)** Anti-serotonin immunoreactive cells, indicated by arrows, can be detected posterior to the pharyngeal area along the nematode body. Confocal images are shown as maximum fluorescence projections. Calibration bar is 20 μm.

**Figure 2.**
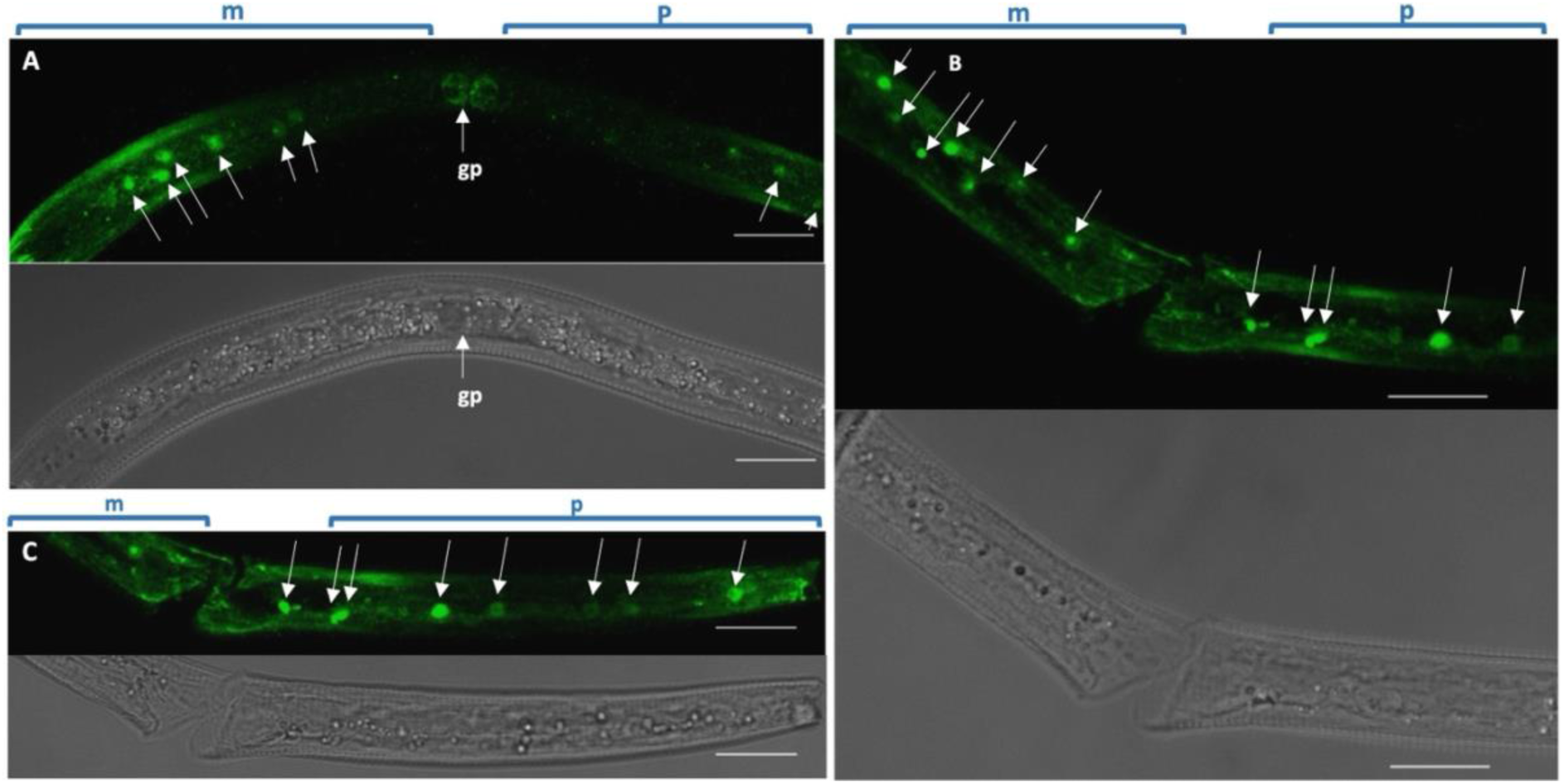
Serotonin immunolocalization in the middle and posterior region of 2nd-stage juveniles (J2s) of *G. pallida*. Anterior region of the J2 nematode is to the left in all images**. (A, B, C)** Anti-serotonin staining is detected in cells indicated by arrows along the nematode body. In **(A)** two large, faintly serotonin-immunoreactive cells are likely to be the genital primordium. Confocal images are shown as maximum fluorescence projections. Middle region of the nematode (m). Posterior region of the nematode (p). Calibration bar is 20 μm.

Posterior to the putative ADF cell bodies is at least one pair, F1,2, and three single cell bodies, G, H and I (Fig. 1B). One unpaired cell body, G, is moderately stained whilst the F1,2, H, and I putative cell bodies are infrequently (30% n=10) and faintly stained. In *C. elegans* posterior to the ADF neurons, the unpaired RIH and I5 neurons and one pair of AIM neurons are serotonin-IR (Serrano-Saiz et al., 2017). However, the fifth putative cell body identified in *G. pallida* has no positional homologue in *C. elegans.* None of these cell bodies have a counterpart in *P. penetrans* and only a putative cell body homologue of the RIH neuron was serotonin-IR in this region in *M. graminicola*. The radial muscles of the median bulb were infrequently (10%, n=10) fluorescent (Fig. 1C).

Puncta of sixteen serotonin-IR cells were also detected along the nematode body (Fig. 1D, Fig.2, Supp Fig. 2, Supp Fig. 3). DAPI staining of nuclei confirmed that these were cells not axonal varicosities associated with neuronal projections (Supp. Fig. 2). At least eight serotonin-IR cells were detected in the middle region of the nematode (Supp. Fig. 2A, B) and a maximum of seven serotonin-IR cells were detected in the posterior region of the nematode (Supp. Fig. 3). Fluorescence was infrequently detected (10%, n=10) in a cell localized in the genital primordium area (Supp. Fig. 2C). None of these serotonin-IR cells are located on the ventral cord (Supp. Fig. 2), therefore we exclude the possibility that they are homologues of the *C. elegans* CP or VC neurons. A control experiment lacking primary antibody validated the observed fluorescence as specific for serotonin-IR cells or structures (Supp. Fig. 4).

**Figure 3.**
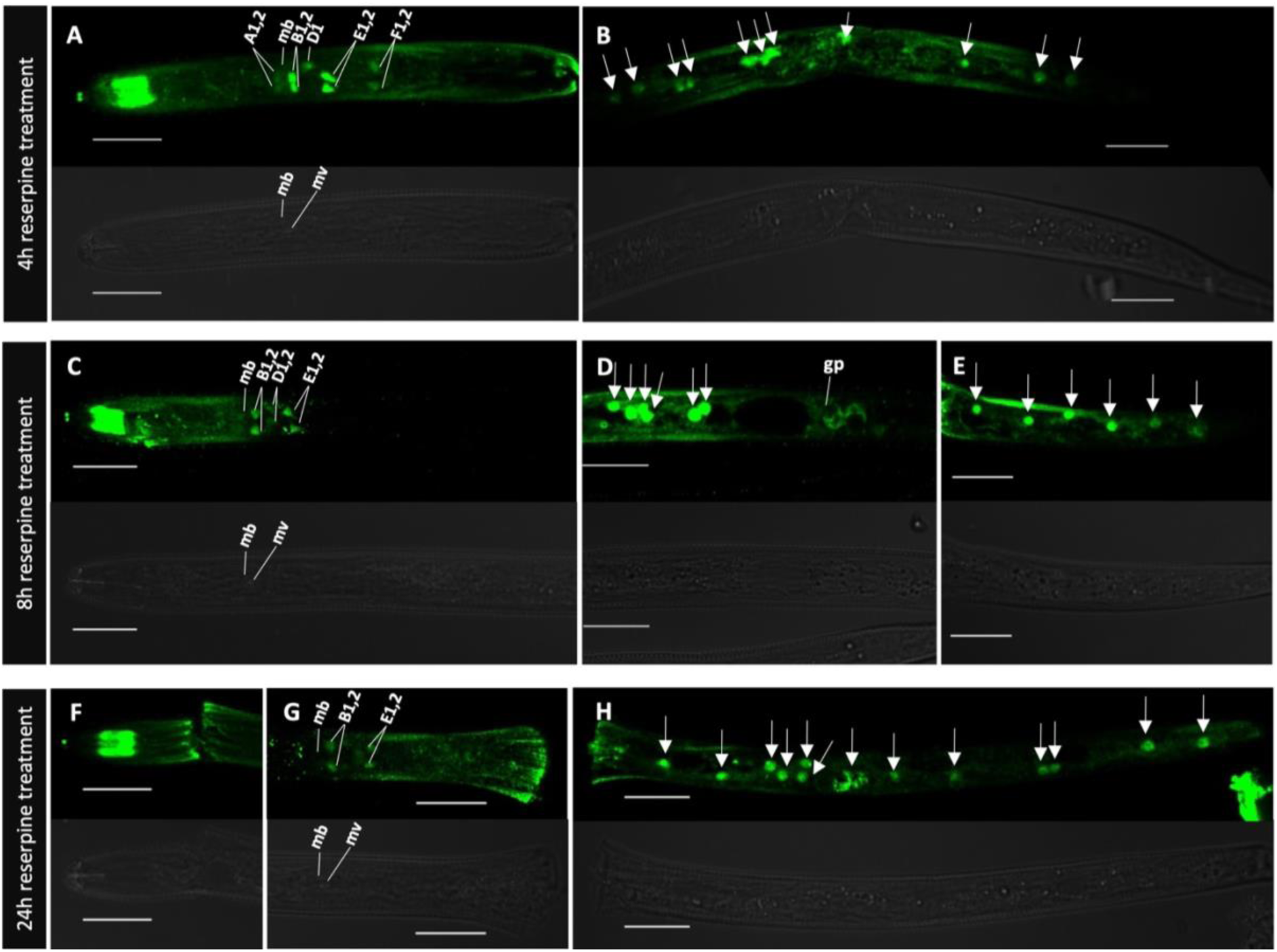
Serotonin immunolocalization in 2nd-stage juveniles (J2s) of *G. pallida* following 4, 8 and 24 hours of reserpine treatment. Anterior region of the *G. pallida* J2 is to the left in all images. **(A, C, F)** No change in the serotonin staining of the amphid pores and pouches is observed even after 24 hours of reserpine treatment. **(A)** After 4 hours of reserpine treatment there are small but visible changes in the pharyngeal area. B1,2 and E1,2 cell bodies, putative homologues of NSM and ADF cell bodies from *C. elegans*, become strongly stained while the other cell bodies are weakly stained in the pharyngeal area. **(B)** No visible changes in the intensity of staining can be observed in the serotonin-IR cells along the body after 4 hours of reserpine treatment. **(C)** After 8 hours of reserpine treatment the changes in the pharyngeal area were similar to those seen after 4 hours of treatment. However, the B1,2 cell bodies were slightly less stained. **(D)** After 8 hours of reserpine treatment the serotonin-IR cells along the nematode body were very strongly stained. **(G)** After 24 hours of reserpine treatment the B1,2 and E1,2 cell bodies were only weakly stained and the other cell bodies in the area, including A1,2 were either occasionally very weakly stained or no longer visible. **(H)** The serotonin-IR cells along the nematode body maintained the pattern described in Fig.2 and they were strongly stained after 24 hours of treatment. Confocal images are shown as maximum fluorescence projections. Calibration bar is 20μm.

**Figure 4.**
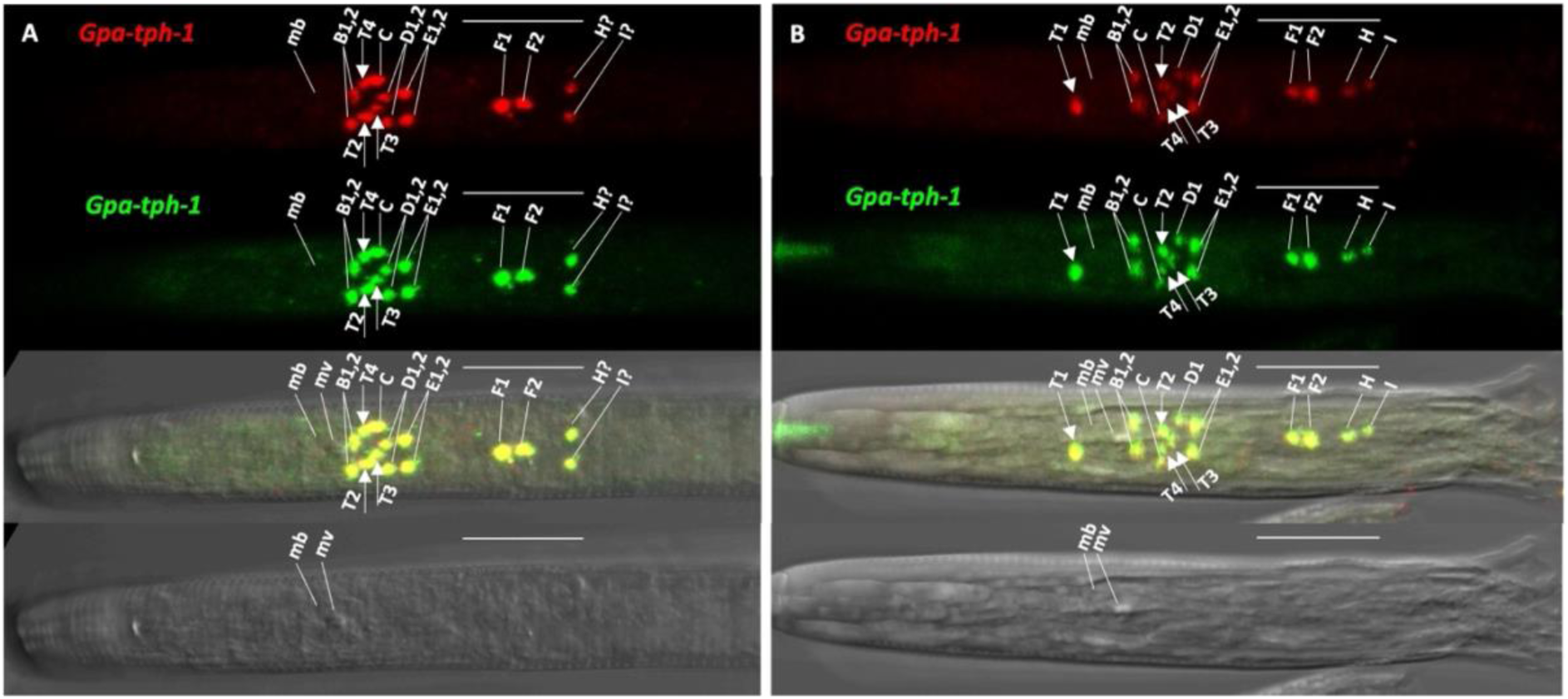
Expression pattern of the *Gpa-tph-1* gene in *G. pallida* J2s using *in situ* HCR. Anterior region of the *G. pallida* J2 is to the left in all images. Two non-overlapping sets of probes, each consisting of fourteen probes were amplified by hairpins labelled with Alexa 488 (green) and Alexa 647 (red) respectively. **(A)** One pair of symmetrically bilateral cell bodies (B1,2 - likely homologues of *C. elegans* NSM neurons) was identified in the pharyngeal median bulb (mb) posterior to the median bulb valve (mv). The putative cell bodies C and D1,2 most likely represent the serotonin-IR cell bodies labelled with the same letters in Fig. 1. Another three putative cell bodies, T2, T3 and T4, marked with arrows are in this region. **(A, B)** The symmetrically bilateral cell bodies E 1,2 are most likely homologues of the ADF neurons in *C. elegans.* Four putative unpaired cell bodies likely matching F, G, H, I from Fig.1 were identified posterior to E 1,2. **(B)**. Infrequently an unpaired cell body, T1, marked with an arrow, was identified anterior to the pharyngeal median bulb. Each panel consists of the Alexa 647 (red) and Alexa 488 (green) fluorescent images, the overlay of these green and red probe sets superimposed on the brightfield image and the brightfield image alone. Confocal images are shown as maximum fluorescence projections. Calibration bar is 20 μm.

### Reserpine treatment induces rapid changes in the serotonin patterning of *G. pallida*

Reserpine potently inhibits the vesicular monoamine transporter (VMAT) preventing the storage of serotonin in the pre-synaptic vesicles and resulting in neuronal depletion (Crisford et al., 2020). We treated *G. pallida* J2s with reserpine for 4, 8 and 24 hours prior to serotonin immunolocalization and monitored changes in the pattern of serotonin immunostaining. Reserpine treatment did not induce any visible changes in the immunoreactivity of amphid pores and pouches at any time points (Fig. 3A, C, F). After 4 hours of reserpine treatment the putative NSM (B1,2) and ADF (E1,2) neurons were strongly reactive while the other putative cell bodies in the anterior region of the nematode were weakly fluorescent (Fig. 3A). This general pattern persisted after 8 hours of reserpine treatment (Fig. 3 C) although many nematodes displayed weaker reactivity of the putative NSM (B1,B2) cell bodies after this time. After 24 hours the putative NSM and ADF cell bodies were both weakly fluorescent (Fig. 3 G) and the other cell bodies in the pharyngeal area were in most cases not visible (Fig. 3G). In contrast, the serotonin-IR cells in the middle and posterior regions of the nematode remained strongly fluorescent even after 24 hours of reserpine treatment (Fig. 3B, D, E, H). The DMF vehicle control did not induce any changes in the staining pattern of serotonin-IR cell bodies even after 24h exposure (Supp. Fig. 5), confirming the changes observed were induced by reserpine.

**Fig. 5.**
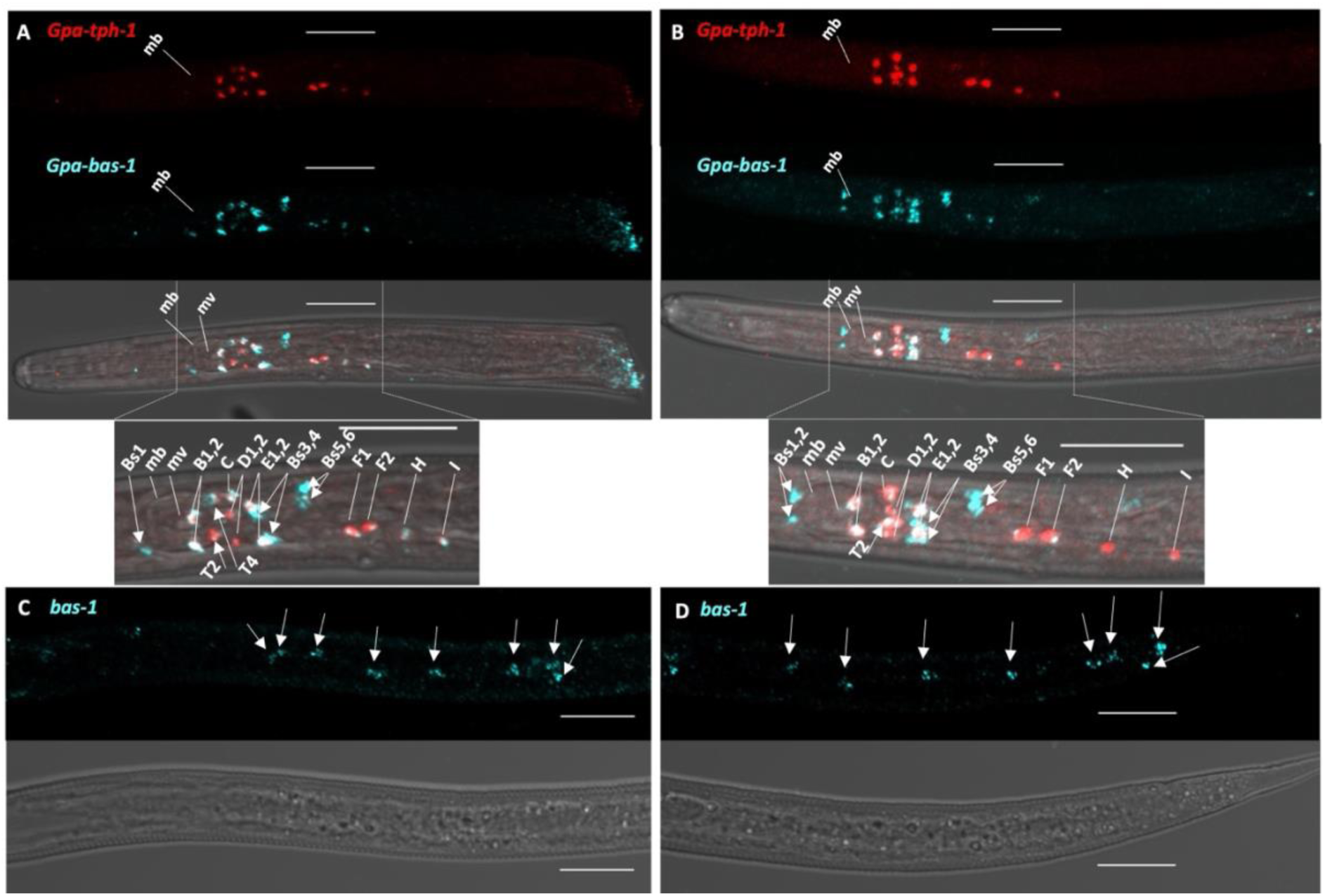
Co-expression pattern of *Gpa-tph-1* and *Gpa-bas-1* genes in *G. pallida* J2s using multiplex *in situ* HCR. Anterior region of the *G. pallida* J2 is to the left in all images. One set of twenty probes targeting *Gpa-bas-1* and one set of twenty-eight probes targeting *Gpa-tph-1,* and amplified with corresponding hairpins labelled with Alexa 546 (blue) and Alexa 647 (red) respectively, were used in a multiplex *in situ* HCR. **(A)** At least thirteen putative cell bodies (B1,2, C, D1,2, E1,2, F1, F2, H, I, T2 and T4) co-express *Gpa-tph-1* and *Gpa-bas-1*. **(B)** Two putative symmetrically bilateral putative cell bodies, Bs1,2, expressing *Gpa-bas-1* only and located anterior to the pharyngeal median bulb (mb), have no counterpart in *C. elegans.* Two additional pairs of putative cell bodies (Bs3,4 and Bs5,6) posterior to the median bulb express *Gpa-bas1* only. One pair (Bs3,4) could be homologues of *C. elegans* CPE while Bs5,6 could be homologues of the ADE neurons. **(C,D)** *Gp-bas-1* was expressed in at least sixteen cells, marked by arrows, in the middle and posterior regions of the nematode. Panels A and B consist of the Alexa 647 (red) and Alexa 546 (blue) fluorescent images, the overlay of these red and blue probe sets superimposed on the brightfield image and the brightfield image alone. Confocal images are shown as maximum fluorescence projections. Calibration bar is 20 μm.

### The *Gpa-tph-1* gene is expressed in fifteen candidate neurons in the pharyngeal region of the nematode

The *tph-1* gene, which is used as a biomarker for serotonergic neurons, encodes tryptophan hydroxylase that catalyses the first step in the biosynthesis of serotonin. The *in situ* hybridization chain reaction method (Sperling and Eves-van den Akker, 2023) was used to analyse the spatial expression of *Gpa-tph-1* in *G. pallida* J2s. This method has high signal-to-noise ratio and its sensitivity can be tuned to enable the detection of genes with relatively low expression level. To test the accuracy of this method in detecting the expression of genes in neurons, two sets of fourteen non overlapping probes were used, each set amplifiable with harpins labelled with different fluorophores. *Gpa*-*tph-1* expression was identified in a total of fifteen cell bodies (Fig. 4). The fluorescence pattern for the two sets of probes overlapped perfectly in all experiments, showing that *in situ* HCR is highly sensitive and effective in detecting gene expression in individual neuronal cell bodies in *G. pallida* (Fig. 4). In the pharyngeal median bulb, posterior to the valve plates, *Gpa-tph-1* is expressed in a pair of symmetrically bilateral cell bodies, B1,2 which most likely are the NSM homologues. Posterior to those there is a cluster of at least eight cell bodies expressing *Gpa-tph-1* (Fig. 4A). Two pairs of symmetrically bilateral cell bodies, D1,2 and E1,2, are easily identifiable alongside another four cell bodies (Fig. 4A). The putative C and D1,2 most likely correspond to their counterparts from the serotonin immunolocalization assay while the putative E1,2 cell bodies are most likely the homologues of the ADF neurons from *C. elegans*. The additional three putative cell bodies, (T2, T3, T4) identified in this area do not have counterparts in our serotonin immunolocalization assay. Infrequently (1%, n=50) during the assays, *Gpa-tph-1* transcripts were detected in one unpaired cell body, T1, anterior to the median bulb (Fig. 4B). Posterior to this cluster, *Gpa-tph-1* is expressed in four cell bodies that might correspond to any of F1,2, G, H, I cell bodies identified in the serotonin immunolocalization assays (Fig. 4). Except for the putative homologues of the NSM (B1,2) and ADF (E1,2) neurons the other cell bodies, eleven in total, have no serotonergic counterpart in *C. elegans.* Expression of *Gpa-tph-1* was not detected along the nematode body.

### *Gpa-bas-1* is co-expressed with *Gpa-tph-1* in at least thirteen neurons in the pharyngeal area of *G. pallida*

The *bas-1* gene encodes an aromatic L-amino acid decarboxylase (AADC) that catalyses the second step in the biosynthesis of serotonin. AADC has a relatively broad range of substrates and is also required for the biosynthesis of dopamine (Hare and Loer, 2004). One set of 20 probes amplified with hairpins labelled with Alexa 546 targeting *Gpa-bas-1* transcripts was used in a multiplex *in situ* HCR alongside one set of 28 probes amplified with hairpins labelled with Alexa 647 targeting *Gpa-tph-1*. This multiplexed experiment identified the same panel of cells revealed by single transcript analysis (Supp. Fig. 6). Thirteen of the fifteen *Gpa-tph-1* expressing cell bodies co-express *Gpa-bas-1* (Fig. 5). A pair of symmetrically bilateral cell bodies, Bs1,2 expressing *Gpa-bas-1* is located anterior to the pharyngeal median bulb (Fig. 5B). These cell bodies do not have a counterpart in *C. elegans.* Inconsistent expression of *Gpa-tph-1* in a cell body in a similar position (Fig. 4B) prevented confirmation that the two genes are co-expressed or not.

**Figure 6.**
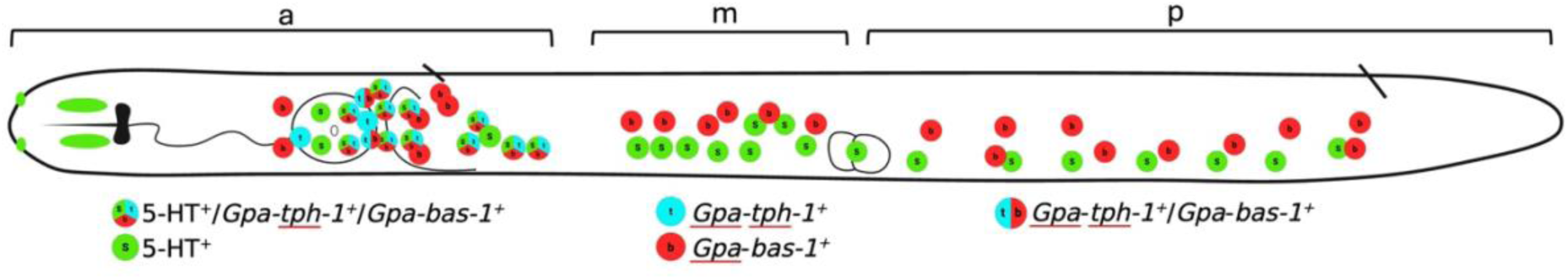
Schematic representation of serotonin immunolocalization and gene expression data. The schematic shows combined serotonin (5-HT) immunoreactivity distribution and expression patterns of *Gpa*-*tph-1* and *Gpa-bas-1* in a *G. pallida J2*. a – anterior region m-middle region, p-posterior region.

At least two cell bodies, Bs3,4 adjacent to the putative ADF homologues (E1,2) express *Gpa-bas-1* only, suggesting they might be homologues of the CEP dopaminergic neurons from *C. elegans.* Posterior to these there is an additional pair of putative cell bodies, Bs5,6 expressing *Gpa-bas-1* only (Fig. 5A,B) suggesting these might be homologues of the ADE dopaminergic neurons from *C. elegans*. *Gpa-bas-1* was expressed along the nematode body in at least sixteen cells in the middle and posterior regions of the nematode (Fig. 5). The *bas-1* gene is expressed in epidermal and intestinal cells in *C. elegans*, as well as PDE neurons in this region so there is a high probability that these cells are of similar type.

## Discussion

Serotonin is an evolutionarily conserved neurotransmitter that plays a central role in regulating nematode behaviour. In the free-living nematode *C. elegans* serotonergic signalling has been extensively characterized and shown to modulate feeding, locomotion, learning, mating, egg laying and other behaviours associated with environmental adaptation and resource exploitation (Sze et al., 2000; Sawin et al., 2000; Shyn et al., 2003; Carnell et al., 2005; Xiao et al., 2006; Bentley et al., 2016). This knowledge has provided an important framework for investigating serotonergic signalling in plant-parasitic nematodes; however, substantial differences in morphology, behaviour and lifestyle, particularly those associated with parasitism and stylet-mediated host invasion, limit direct extrapolation from *C. elegans* (Coke et al., 2024). In contrast to its predominantly modulatory role in free-living and animal-parasitic nematodes, serotonin is a critical regulator of parasitic behaviours in plant-parasitic nematodes, controlling processes required for host invasion and feeding site establishment (Crisford et al., 2020; Fleming et al., 2017). Despite this importance, the serotonergic nervous system of plant-parasitic nematodes remains poorly defined. Here, by comprehensively characterizing the serotonergic system of *Globodera pallida* J2s, we provide a foundation for understanding how serotonin regulates behaviours essential for parasitism and identify key similarities and differences with the serotonergic circuitry described in *C. elegans* and other nematode species.

We mapped serotonin-like immunoreactivity and the expression of the two genes encoding the enzymes required for serotonin synthesis with the aim of identifying serotonergic neurons in the *G. pallida* J2 infective stage. Serotonergic neurons should be positive for all three labelling approaches. Moreover, the serotonin-like immunoreactivity in those neurons should be depleted by reserpine treatment as this will prevent serotonergic neurons storing the neurotransmitter. Based on these criteria, we can confidently predict at least eleven serotonergic neurons in *G. pallida* (Table 1, Fig. 6). Apart from the putative NSM and ADF homologues there are an additional seven candidate serotonergic neurons in *G. pallida* compared to *C. elegans*. Based on their localization five of these could be the homologues of *C. elegans* neurons that only uptake serotonin while two are unknown, suggesting a functional shift in neurotransmitter phenotype and an increased complexity of serotonin circuitry in *G. pallida*. An additional two cell bodies expressing both serotonin synthesising genes were not serotonin-IR, suggesting serotonin levels might have been below our detection threshold or these cells do not accumulate serotonin. Exposure of J2s to reserpine produced visible changes in all these putative cell bodies in the pharyngeal area with a complete loss in the serotonin immunoreactive putative cell bodies A1,2 after 24 hours of exposure, suggesting these are serotonin up-taking cell bodies, as they also do not express *tph-1* and *bas-1*. Differences in staining intensity between different putative serotonergic neurons after reserpine exposure suggest that serotonin might be synthesised at different levels, with the highest in the putative NSM and ADF homologues.

**Table 1.**
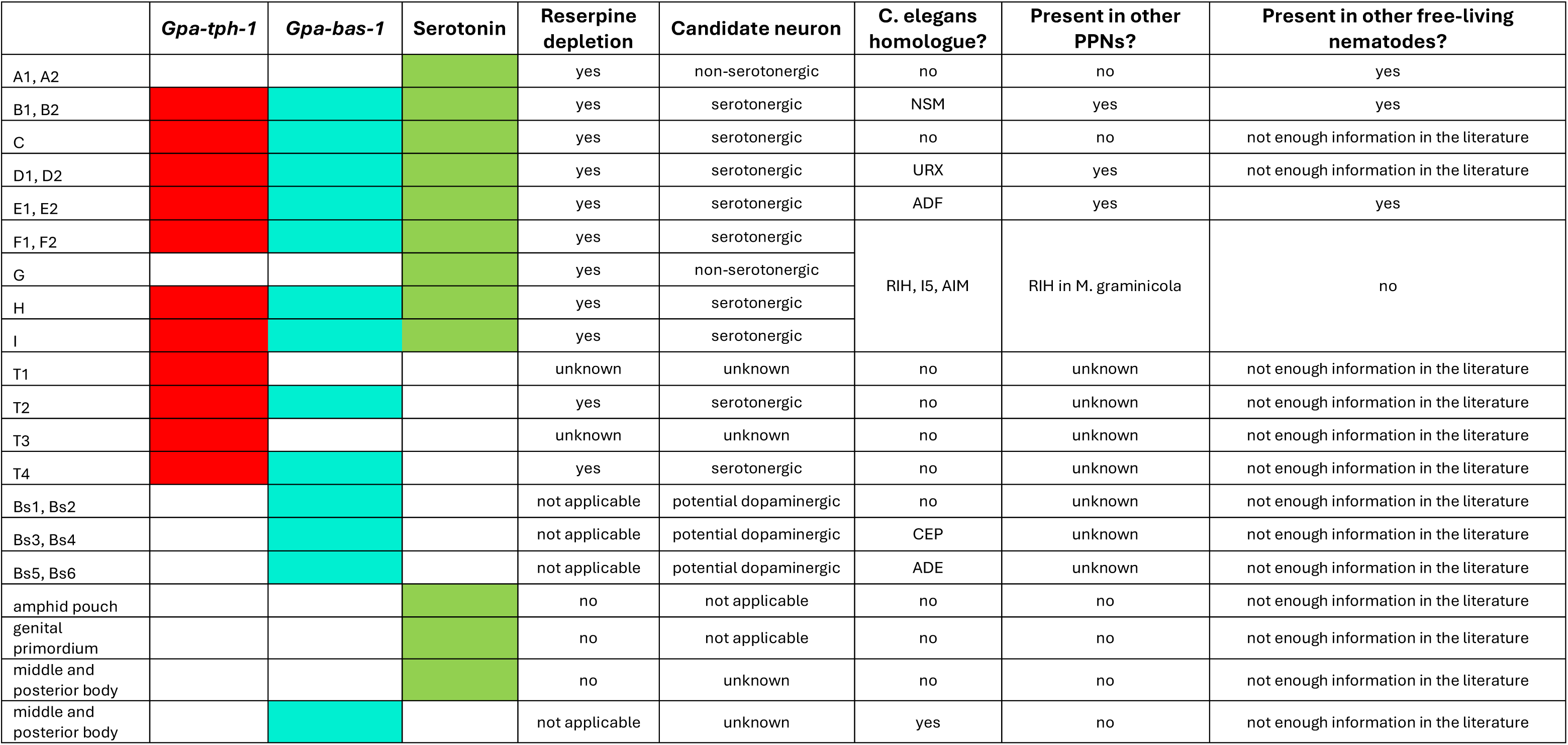
Summary of the serotonin immunolocalization and *Gpa-tph-1* and *Gpa-bas-1* gene expression data.

Fourteen serotonin-IR putative cell bodies detected in the pharyngeal area of *G. pallida* J2s (Table 1, Fig. 6) is twice the number reported in *M. graminicola* (Yang et al., 2024). Despite the fact that the number of serotonergic neurons in other plant parasitic nematodes is unknown, this variation alone suggests a high plasticity of the serotonin system. A similar feature was previously reported in free-living nematodes, where differences in serotonin distribution were linked to the evolutionary adaptation of these nematodes to diverse habitats (Rivard et al., 2010). Conversely, the reduced number of serotonin-IR neurons in the nervous system of animal parasitic nematodes compared to *C. elegans* was associated with their shielded lifestyle, in a stable and predictable environment within the host (Johnson et al., 1996; Rao et al., 2011).

A pair of cell bodies reported as serotonin-IR in both root-knot and root-lesion nematodes (Han et al., 2017; Yang et al., 2024) may represent homologues of the putative URX neurons identified as serotonergic in this study. In *C. elegans* these are sensory neurons with a critical role in oxygen detection, lipid metabolism and lifespan regulation. Their function should be essential for plant parasitic infective stages navigating through the soil towards the root and integrating different cues to find a host. More studies are required to understand how serotonin affects these functions in plant parasitic nematodes. We identified two putative cell bodies that express *Gpa-tph-1* only, suggesting a potential physiological role for TPH-1 in *G. pallida* beyond its role in serotonin synthesis. There are similar reports of *tph-1* expression in *C. elegans* MI, AFD (Wang et al., 2024), AIM and RIH neurons (Sze et al., 2000) even though none of these neurons express *bas-1* (Wang et al., 2024).

Our study also identified putative homologues of the dopaminergic ADE and CEP neurons, and two *Gpa-bas-1* expressing putative cell bodies anterior to the median bulb that have no counterpart in any other nematodes to date. Future work will need to determine if these are truly dopaminergic. An additional sixteen *Gpa-bas-1* expressing cells were identified along the J2 nematode body. Two of these could be homologues of the dopaminergic PDE neurons, while the rest might be epidermal and intestinal cells. The lack of a neuronal atlas in plant-parasitic nematodes makes their precise identification impossible.

We identified three serotonin-IR neurons in the pharyngeal area of *G. pallida* that do not express either of the synthesis genes suggesting these cells most likely only uptake serotonin from the surroundings. They could have a similar function to those in *C. elegans*, as spatio-temporal regulators of serotonin concentration in the pseudocoelomic environment (Jafari et al., 2011). Complementary studies on the expression pattern of the serotonin reuptake transporter MOD-5 in *G. pallida* would be required to confirm this hypothesis. In addition to the canonical neuronal synthesis, serotonin and serotonin-derivate metabolites produced via a parallel pathway in the epidermal cells of *C. elegans,* which co-express phenylalanine transferase and *bas-1*, contribute to the previously described serotonin-dependent phenotypes in *C. elegans* (Yu et al., 2023). We identified sixteen strongly stained serotonin-IR cells along the body of the nematode, and their position, away from the ventral cord, suggests they are non-neuronal cells. The staining of these cells was not affected by the reserpine treatment, suggesting the presence of serotonin derivatives that could accumulate from the pseudocoelomic fluid or are synthesised through a parallel pathway similar to that described in *C. elegans*.

Our immuno-analysis also revealed unexpected serotonin-IR in amphid pouches and pores further suggesting the presence of serotonin derivatives in these structures. Plant-parasitic nematodes rely on chemoreception to locate a host in soil by detecting various environmental cues (Curtis, 2007). The amphids are the main chemosensory organs, containing dendritic processes in an amphid pouch which are bathed in secretions produced by the amphid sheath cells (Perry, 1993). The amphid pouch opens to the external environment though a pore allowing the dendritic processes to interact with the surroundings. The secretions have a role in maintaining the structural and functional integrity of the dendritic endings. *C. elegans* mutants with defects in secretion production display an impaired sensory function. It is not known if serotonin-IR metabolites are produced by the sheath cells, but their accumulation in the amphids is not affected by reserpine treatment, suggesting that their production may involve a distinct serotonin biosynthesis pathway.

Overall, the serotonin-system in *G. pallida* seems more complex than that of *C. elegans* and it is distinct from other plant-parasitic nematodes studied to date. The high number of serotonergic neurons identified in *G. pallida* suggests differences in the synaptic connections and functional properties of putative homologous neurons between cyst nematodes and the model *C. elegans*. These differences highlight substantial divergence in serotonin signalling between the free-living *C. elegans* and the plant-parasitic cyst nematode *G. pallida*, despite broad conservation of nematode nervous system architecture, underscoring the limitations of extrapolating knowledge from establish model organisms to parasitic systems. HCR *in situ* hybridization offer a valuable approach to investigate neuronal organization and signalling in non-model nematode species with limited or unavailable reverse-genetic tools.

## Funding

This work was funded through Biotechnology and Biological Sciences Research Council (BBSRC) grants BB/T001097/1 and BB/T002867/1. R.H. was supported by a PhD studentship from BBSRC White Rose Doctoral Training Partnership (DTP).

## Notes

### Competing Interest Statement

The authors have declared no competing interest.

## References

Bentley, B., Branicky, R., Barnes, C. L., Chew, Y. L., Yemini, E., Bullmore, E. T., Vértes, P. E. & Schafer, W. R. 2016. The Multilayer Connectome of *Caenorhabditis elegans*. PLoS Comput Biol, 12, e1005283.

Calahorro, F., Chapman, M., Dudkiewicz, K., Holden-Dye, L. & O’connor, V. 2022. PharmacoGenetic targeting of a *C. elegans* essential neuron provides an *in vivo* screening for novel modulators of nematode ion channel function. Pestic Biochem Physiol, 186, 105152.

Carnell, L., Illi, J., Hong, S. W. & Mcintire, S. L. 2005. The G-protein-coupled serotonin receptor SER-1 regulates egg laying and male mating behaviors in *Caenorhabditis elegans*. J Neurosci, 25, 10671–81.

Choi, H. M., Calvert, C. R., Husain, N., Huss, D., Barsi, J. C., Deverman, B. E., Hunter, R. C., Kato, M., Lee, S. M., Abelin, A. C., Rosenthal, A. Z., Akbari, O. S., Li, Y., Hay, B. A., Sternberg, P. W., Patterson, P. H., Davidson, E. H., Mazmanian, S. K., Prober, D. A., Van De Rijn, M., Leadbetter, J. R., Newman, D. K., Readhead, C., Bronner, M. E., Wold, B., Lansford, R., Sauka-Spengler, T., Fraser, S. E. & Pierce, N. A. 2016. Mapping a multiplexed zoo of mRNA expression. Development, 143, 3632–3637.

Coke, M. C., Bell, C. A. & Urwin, P. E. 2024. The use of *Caenorhabditis elegans* as a model for plant-parasitic nematodes: What have we learned? Annu Rev Phytopathol, 62, 157–172.

Cotton, J. A., Lilley, C. J., Jones, L. M., Kikuchi, T., Reid, A. J., Thorpe, P., Tsai, I. J., Beasley, H., Blok, V., Cock, P. J., Eves-Van Den Akker, S., Holroyd, N., Hunt, M., Mantelin, S., Naghra, H., Pain, A., Palomares-Rius, J. E., Zarowiecki, M., Berriman, M., Jones, J. T. & Urwin, P. E. 2014. The genome and life-stage specific transcriptomes of *Globodera pallida* elucidate key aspects of plant parasitism by a cyst nematode. Genome Biol, 15, R43.

Crisford, A., Calahorro, F., Ludlow, E., Marvin, J. M. C., Hibbard, J. K., Lilley, C. J., Kearn, J., Keefe, F., Johnson, P., Harmer, R., Urwin, P. E., O’connor, V. & Holden-Dye, L. 2020. Identification and characterisation of serotonin signalling in the potato cyst nematode *Globodera pallida* reveals new targets for crop protection. PLoS Pathog, 16, e1008884.

Curtis, R. H. C. 2007. Plant parasitic nematode proteins and the host–parasite interaction. Briefings in Functional Genomics & Proteomics, 6, 50–58.

Desaeger, J., Wram, C. & Zasada, I. 2020. New reduced-risk agricultural nematicides - rationale and review. Journal of Nematology, 52, 1–16.

Erickson, J. D., Eiden, L. E. & Hoffman, B. J. 1992. Expression cloning of a reserpine-sensitive vesicular monoamine transporter. Proc Natl Acad Sci U S A, 89, 10993–7.

Fleming, T. R., Maule, A. G. & Fleming, C. C. 2017. Chemosensory responses of plant parasitic nematodes to selected phytochemicals reveal long-term habituation traits. J Nematol, 49, 462–471.

Han, J. & Schroeder, N. E. 2025. Neuronal basis of host-finding and feeding in plant-parasitic nematodes. Annual Review of Phytopathology, 63, 383–402.

Han, Z., Boas, S. & Schroeder, N. E. 2017. Serotonin regulates the feeding and reproductive behaviors of *Pratylenchus penetrans*. Phytopathology, 107, 872–877.

Hare, E. E. & Loer, C. M. 2004. Function and evolution of the serotonin-synthetic *bas-1* gene and other aromatic amino acid decarboxylase genes in *Caenorhabditis*. BMC Evolutionary Biology, 4, 24.

Holden-Dye, L. & Walker, R. J. 2014. Anthelmintic drugs and nematicides: studies in *Caenorhabditis elegans*. WormBook, 1-29.

Hong, R. L., Riebesell, M., Bumbarger, D. J., Cook, S. J., Carstensen, H. R., Sarpolaki, T., Cochella, L., Castrejon, J., Moreno, E., Sieriebriennikov, B., Hobert, O. & Sommer, R. J. 2019. Evolution of neuronal anatomy and circuitry in two highly divergent nematode species. eLife, 8, e47155.

Horvitz, H. R., Chalfie, M., Trent, C., Sulston, J. E. & Evans, P. D. 1982. Serotonin and octopamine in the nematode *Caenorhabditis elegans*. Science, 216, 1012–4.

Jafari, G., Xie, Y., Kullyev, A., Liang, B. & Sze, J. Y. 2011. Regulation of extrasynaptic 5-HT by serotonin reuptake transporter function in 5-HT-absorbing neurons underscores adaptation behavior in *Caenorhabditis elegans*. J Neurosci, 31, 8948–57.

Johnson, C. D., Reinitz, C. A., Sithigorngul, P. & Stretton, A. O. 1996. Neuronal localization of serotonin in the nematode *Ascaris suum*. J Comp Neurol, 367, 352–60.

Kearn, J., Ludlow, E., Dillon, J., O’connor, V. & Holden-Dye, L. 2014. Fluensulfone is a nematicide with a mode of action distinct from anticholinesterases and macrocyclic lactones. Pesticide Biochemistry and Physiology, 109, 44–57.

Kimber, M. J., Fleming, C. C., Prior, A., Jones, J. T., Halton, D. W. & Maule, A. G. 2002. Localisation of *Globodera pallida* FMRFamide-related peptide encoding genes using in situ hybridisation. International Journal for Parasitology, 32, 1095–1105.

Lilley, C. J., Atkinson, H. J. & Urwin, P. E. 2005. Molecular aspects of cyst nematodes. Mol Plant Pathol, 6, 577–88.

Loer, C. M. & Kenyon, C. J. 1993. Serotonin-deficient mutants and male mating behavior in the nematode *Caenorhabditis elegans*. J Neurosci, 13, 5407–17.

Masler, E. 2008. Responses of *Heterodera glycines* and *Meloidogyne incognita* to exogenously applied biogenic amines. Nematology, 10, 911–917.

Rao, V. T., Forrester, S. G., Keller, K. & Prichard, R. K. 2011. Localisation of serotonin and dopamine in *Haemonchus contortus*. Int J Parasitol, 41, 249–54.

Rivard, L., Srinivasan, J., Stone, A., Ochoa, S., Sternberg, P. W. & Loer, C. M. 2010. A comparison of experience-dependent locomotory behaviors and biogenic amine neurons in nematode relatives of *Caenorhabditis elegans*. BMC Neuroscience, 11, 22.

Rodriguez Araujo, N., Hernando, G., Corradi, J. & Bouzat, C. 2022. The nematode serotonin-gated chloride channel MOD-1: A novel target for anthelmintic therapy. Journal of Biological Chemistry, 298.

Sawin, E. R., Ranganathan, R. & Horvitz, H. R. 2000. *C. elegans* locomotory rate is modulated by the environment through a dopaminergic pathway and by experience through a serotonergic pathway. Neuron, 26, 619–31.

Schleker, A. S. S., Rist, M., Matera, C., Damijonaitis, A., Collienne, U., Matsuoka, K., Habash, S. S., Twelker, K., Gutbrod, O., Saalwächter, C., Windau, M., Matthiesen, S., Stefanovska, T., Scharwey, M., Marx, M. T., Geibel, S. & Grundler, F. M. W. 2022. Mode of action of fluopyram in plant-parasitic nematodes. Scientific Reports, 12, 11954.

Serrano-Saiz, E., Pereira, L., Gendrel, M., Aghayeva, U., Bhattacharya, A., Howell, K., Garcia, L. R. & Hobert, O. 2017. A neurotransmitter atlas of the *Caenorhabditis* elegans male nervous system reveals sexually dimorphic neurotransmitter usage. Genetics, 206, 1251–1269.

Shyn, S. I., Kerr, R. & Schafer, W. R. 2003. Serotonin and Go modulate functional states of neurons and muscles controlling *C. elegans* egg-laying behavior. Curr Biol, 13, 1910–5.

Sobczak, M. & Golinowski, W. 2011. Cyst Nematodes and Syncytia. In: Jones, J., Gheysen, G. & Fenoll, C. (eds.) Genomics and Molecular Genetics of Plant-Nematode Interactions. Dordrecht: Springer Netherlands.

Sperling, A. L. & Eves-Van Den Akker, S. 2023. Whole mount multiplexed visualization of Dna, mrna, and protein in plant-parasitic nematodes. Plant Methods, 19, 139.

Sze, J. Y., Victor, M., Loer, C., Shi, Y. & Ruvkun, G. 2000. Food and metabolic signalling defects in a *Caenorhabditis elegans* serotonin-synthesis mutant. Nature, 403, 560–4.

Thorpe, P., Mantelin, S., Cock, P. J., Blok, V. C., Coke, M. C., Eves-Van Den Akker, S., Guzeeva, E., Lilley, C. J., Smant, G., Reid, A. J., Wright, K. M., Urwin, P. E. & Jones, J. T. 2014. Genomic characterisation of the effector complement of the potato cyst nematode *Globodera pallida*. BMC Genomics, 15, 923.

Trudgill, D. L., Phillips, M. S. & Elliott, M. J. 2014. Dynamics and management of the white potato cyst nematode *Globodera pallida* in commercial potato crops. Annals of Applied Biology, 164, 18–34.

Wang, C., Vidal, B., Sural, S., Loer, C., Aguilar, G. R., Merritt, D. M., Toker, I. A., Vogt, M. C., Cros, C. & Hobert, O. 2024. A neurotransmitter atlas of *C. elegans* males and hermaphrodites. eLife Sciences Publications, Ltd.

Xiao, H., Hapiak, V. M., Smith, K. A., Lin, L., Hobson, R. J., Plenefisch, J. & Komuniecki, R. 2006. SER-1, a *Caenorhabditis elegans* 5-HT2-like receptor, and a multi-PDZ domain containing protein (MPZ-1) interact in vulval muscle to facilitate serotonin-stimulated egg-laying. Developmental Biology, 298, 379–391.

Yang, Z., Zhang, Z., Long, X., Shi, X., Wang, D., Peng, D., Ye, S. & Ding, Z. 2024. Clarifying the functional role of serotonin in *Meloidogyne graminicola* host plant parasitism by immunolocalization and RNA interference. Phytopathology, 114, 1401–1410.

Yemini, E., Lin, A., Nejatbakhsh, A., Varol, E., Sun, R., Mena, G. E., Samuel, A. D. T., Paninski, L., Venkatachalam, V. & Hobert, O. 2021. NeuroPAL: A multicolor atlas for whole-brain neuronal identification in *C. elegans*. Cell, 184, 272–288.e11.

Yu, J., Vogt, M. C., Fox, B. W., Wrobel, C. J. J., Fajardo Palomino, D., Curtis, B. J., Zhang, B., Le, H. H., Tauffenberger, A., Hobert, O. & Schroeder, F. C. 2023. Parallel pathways for serotonin biosynthesis and metabolism in *C. elegans*. Nat Chem Biol, 19, 141–150.

