## Supplemental Data for "The potato cyst nematode *Globodera pallida* possesses an expanded serotonergic nervous system"

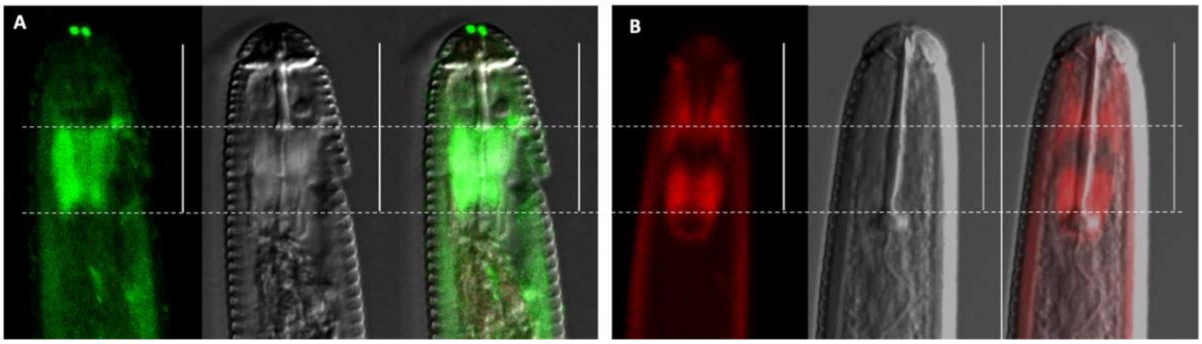


**Supp. Fig. 1. Serotonin immunolocalization and phalloidin staining in the head of *G. pallida* J2s. (A)** The pattern of serotonin immunolocalization in the head of *G. pallida* is not similar to **(B)** the pattern of phalloidin staining (red) of the stylet protractor muscles. Confocal images are shown as maximum fluorescence projections. Calibration bar is 20 μm.

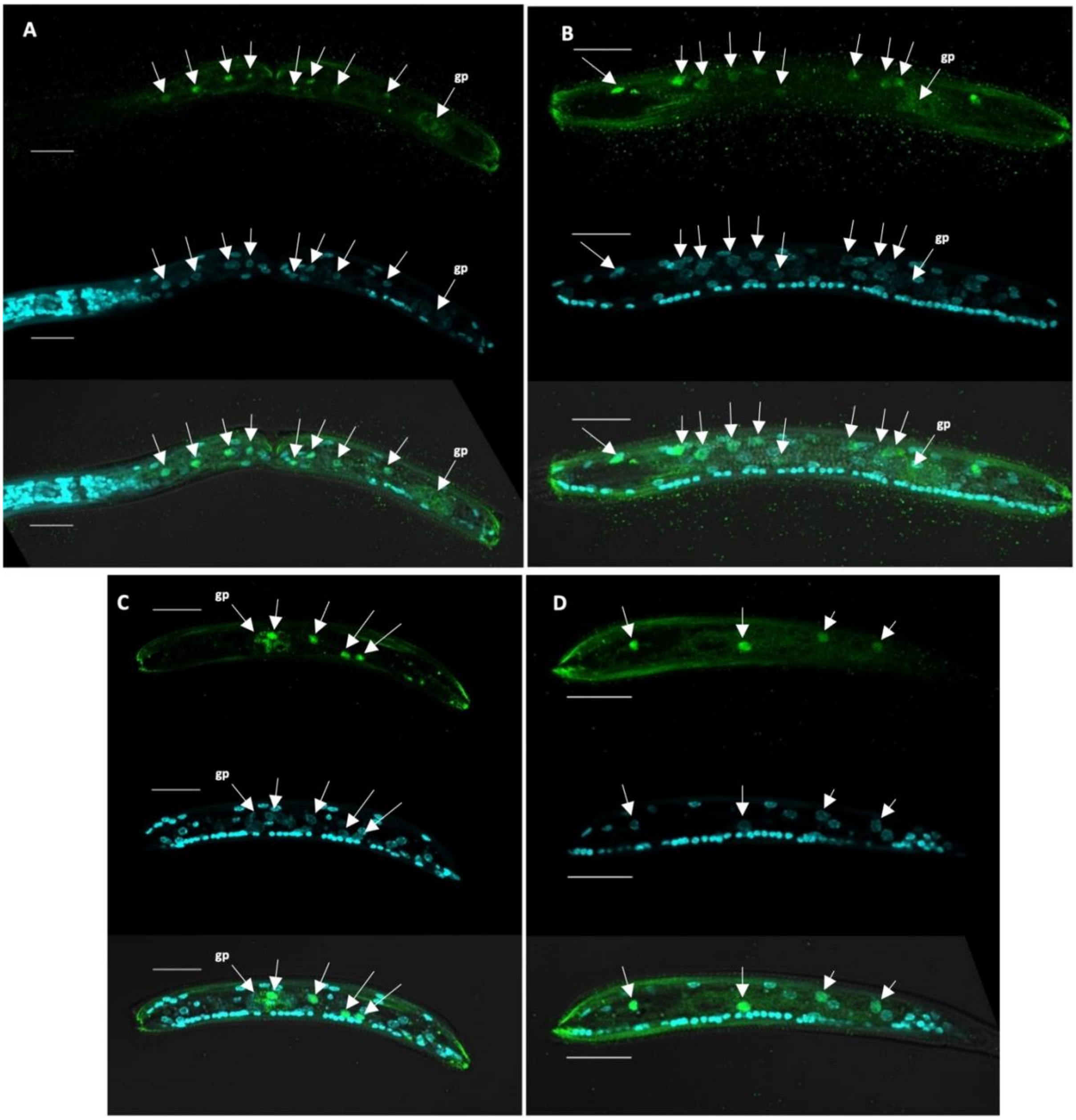


**Supp. Fig. 2. Serotonin immunolocalization and DAPI staining in the middle and posterior regions of *G. pallida* J2s.** Anterior region of *G. pallida* J2 is to the left in A, B and D, unknown in C. **(A & B)** There are at least 9 stained cells anterior to the genital primordium. **(B,C,D)** All serotonin immunoreactive cells in the middle and posterior region of nematode are DAPI (cyan) stained, indicating the presence of a nucleus. None of these stained cells are situated on the ventral cord. **(C)** Infrequently, one cell is stained in the area of the genital primordium. Confocal images are shown as maximum fluorescence projections. Calibration bar is 20 μm.

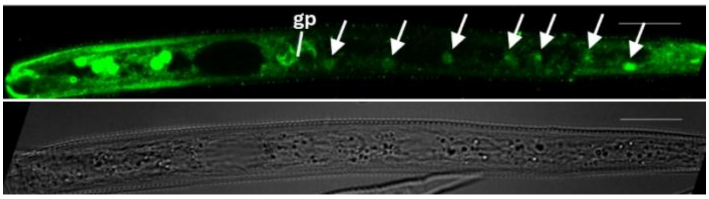


**Supp. Fig. 3**. **Serotonin immunolocalization in the posterior region of *G. pallida* J2.** Anterior region of *G. pallida* J2 is to the left. There are at least seven serotonin immunoreactive cells posterior to the genital primordium (gp). Confocal images are shown as maximum fluorescence projections. Calibration bar is 20 μm.

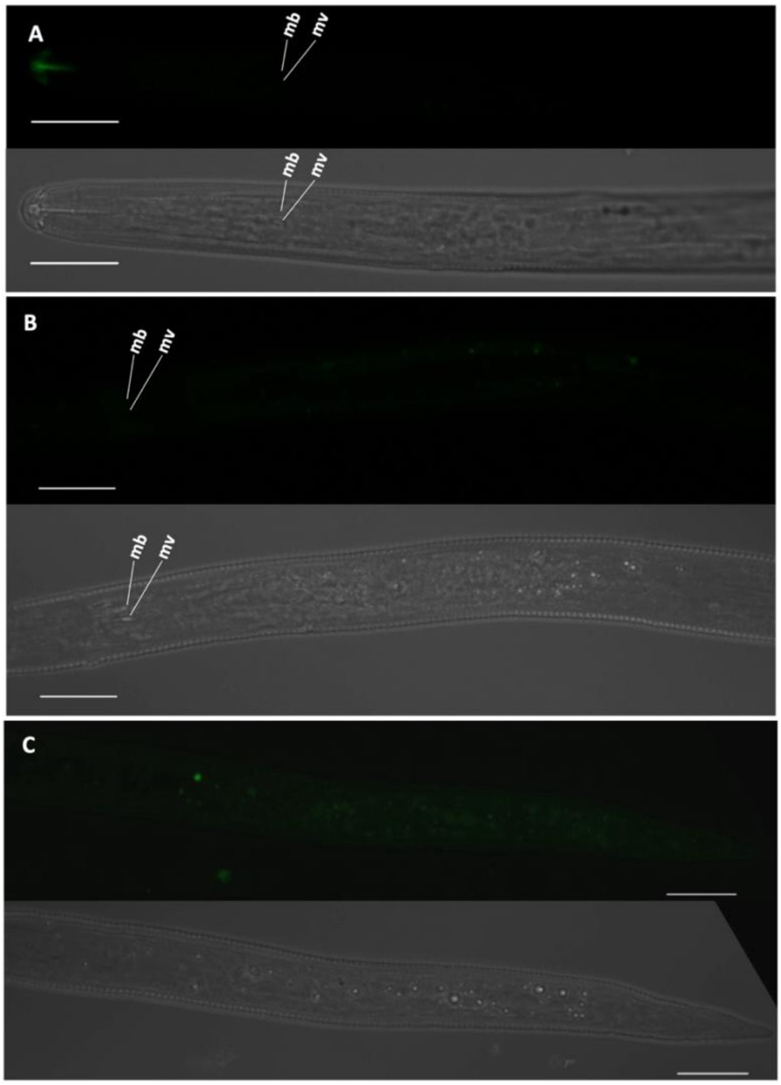


**Supp. Fig. 4. Immunostaining of *G. pallida* J2s in the absence of the primary antibody.** Anterior region of *G. pallida* is to the left in all images. **(A, B ,C)** The secondary antibody does not bind to any cells or structures in the nematode in the absence of the primary antibody. **(A)** Anterior region of the nematode **(B)** Middle region of the nematode **(C)** Posterior region of the nematode. Median bulb (mb); median bulb valve (mv). Confocal images are shown as maximum fluorescence projections. Calibration bar is 20 μm.

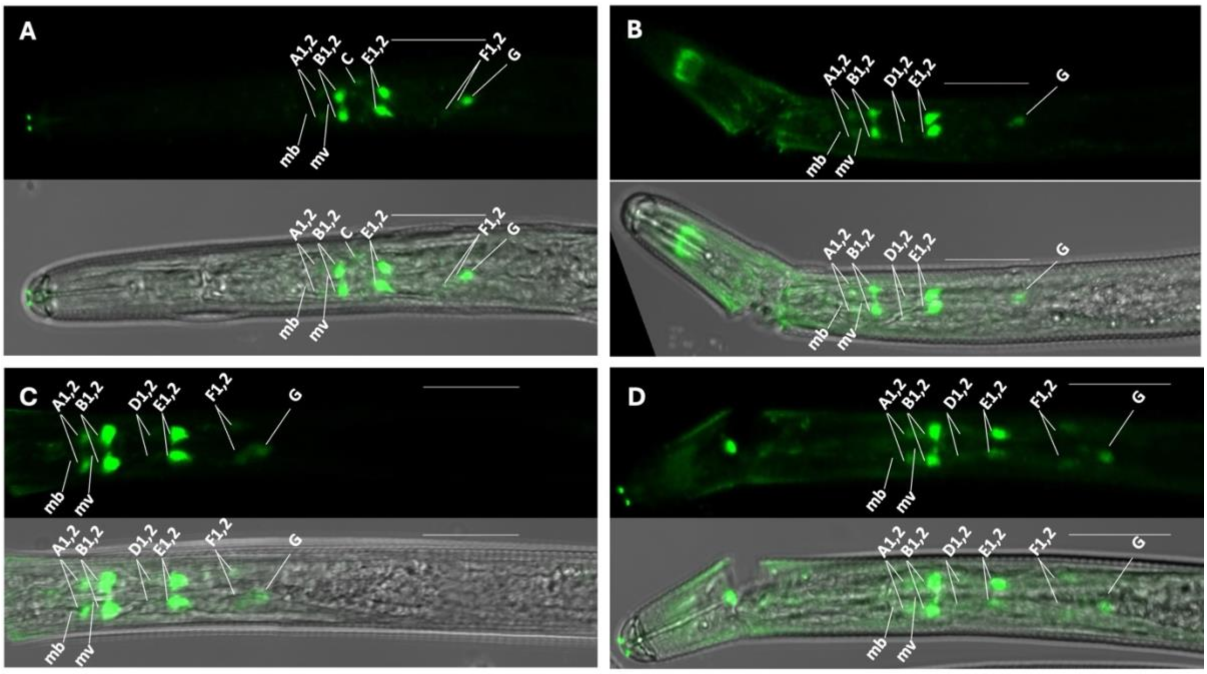


**Supp. Fig. 5. Serotonin immunolocalization after 0 (A), 4 (B), 8 (C) and 24 h (D) incubation in 0.33% DMF (vehicle for reserpine)**. Anterior region of *G. pallida* J2 is to the left in all images. Incubation in 0.33% DMF (vehicle for reserpine) did not induce the changes in serotonin immunoreactivity observed following reserpine treatments. Confocal images are shown as maximum fluorescence projections. Calibration bar is 20 μm.

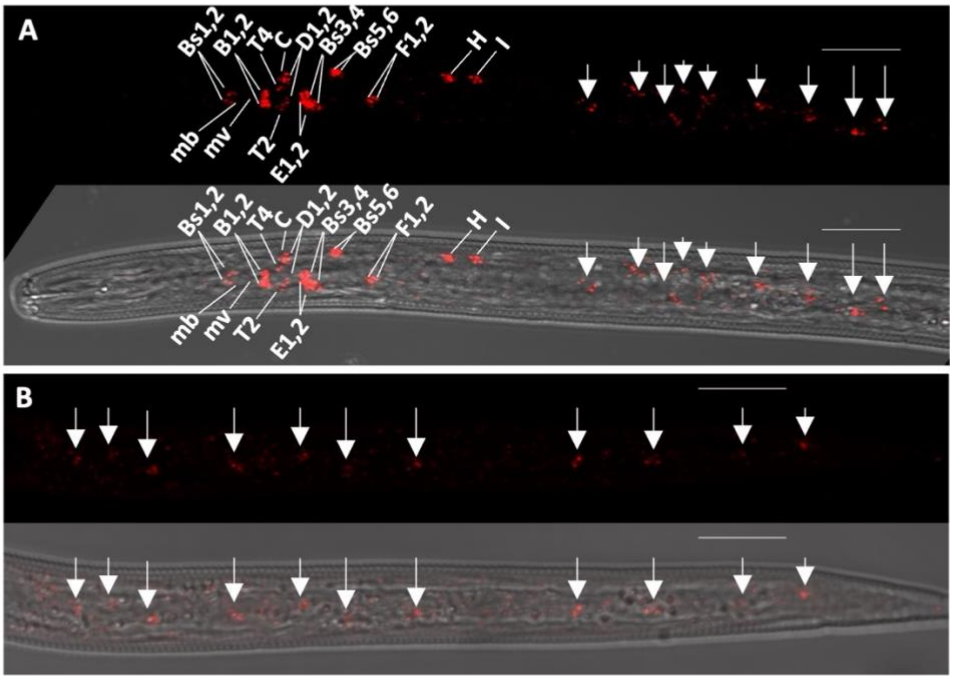


**Supp. Fig. 6. Expression pattern of *Gpa-bas-1* in *G. pallida* J2s using *in situ* HCR**. Anterior region of the *G. pallida* J2 is to the left in all images. One set of twenty probes amplified with hairpins labelled with Alexa 546 were used. **(A)** The expression of *Gpa-bas-1* in the anterior region of the nematode was detected in nineteen cells, a pattern identical to that of *Gpa-bas-1* detected during the multiplex *in situ* HCR. **(B)** *Gpa-bas-1* was expressed in cells marked by arrows in the middle and posterior regions of the nematode. Panels A and B consist of the fluorescent image (Alexa 546 - red) and the fluorescent image superimposed on the brightfield image. Confocal images are shown as maximum fluorescence projections. Calibration bar is 20 μm.
